# PeSTo: parameter-free geometric deep learning for accurate prediction of protein interacting interfaces

**DOI:** 10.1101/2022.05.09.491165

**Authors:** Lucien F. Krapp, Luciano A. Abriata, Fabio Cortés Rodriguez, Matteo Dal Peraro

## Abstract

Predicting the interactions that a protein can establish with other molecules from its structure remains a major challenge. As shown by recent applications to tertiary structure prediction and opposite to current mainstream methods for interaction interface prediction, low-level, geometry-based, physicochemical-agnostic representations of structures have several advantages over methods that require pre-calculation of surfaces, charges, hydrophobicity, and other kinds of parameterizations. Here we introduce a new geometric transformer that acts directly on protein atoms labelled with nothing more than element names. The resulting model outperforms the state of the art for the prediction of protein-protein interaction interfaces and distinguishes interfaces with nucleic acids, lipids, small molecules and ions with high confidence. The low computational cost of this method (available online at https://pesto.epfl.ch/) enables processing high volumes of structural data, such as molecular dynamics trajectories allowing the discovery of interfaces that remain inconspicuous in static experimentally solved structures.

## Introduction

Molecular interfaces are ubiquitous in biology and of utmost relevance beyond their central role in establishing cell boundaries and intracellular organization [1-3]. Especially so around proteins, which perform their functions by interacting with other proteins as well as with nucleic acids, membranes, and small molecules and ions of various kinds. Predicting the interactions that a given protein can establish with other molecules remains a major problem in biology, still open despite numerous developments along various fronts [4-7]. The most modern methods for predicting protein interactions currently target either specific pairs of interacting residues/atoms, relying intensively on residue-residue coevolution, and thus limited to protein-protein interactions, or predicting which regions of a protein are prone to interaction [7-14]. Even the latter, presumably a simpler problem, is yet far from solved, and most methods aim mainly at discovering protein interfaces tailored to interact with other proteins, with a strong focus on features of the protein surface and in some cases also exploiting sequence conservation at the surface. The main limitation of these methods is that calculations of protein surfaces and mapping of their properties are time-consuming, complicating their high-throughput application to proteomes scale, very sensitive to details and errors in the 3D structure or model; while relying on sequence conservation might make the system fail at shallow sequence alignments. Folding protein complexes discover the interaction interface, the structure of the complex, and the conformations of the subunits. However, it is significantly slower than predicting the interaction interface and will fail if the folding itself fails [15]. Here, building on the recent successful application of transformers to various problems in natural language processing and protein structure prediction, we developed a new rotationally invariant transformer-based neural network that acts directly on protein atoms predicting interaction interfaces with high confidence, without the need for parameterization of the system’s physics nor for any surface calculation, running fast enough to process large structural ensembles, such those generated from molecular dynamics. Trained to predict protein-protein interaction interfaces our method outperforms the state of the art. Training to predict other kinds of protein interfaces is straightforward as the method does not depend on explicit description, and thus assignment of physicochemical features.

### The protein structure transformer (PeSTo)

Protein structures can be represented in multiple ways: surfaces describing electrochemical properties [7,16,17], volumes using voxels of densities [18,19], graphs of connected atoms [20], or point clouds. The choice of a representation for a specific problem is important as it will affect the overall performance of a machine learning model [21]. Scalar quantities of proteins such as the energy or interactions interfaces are intrinsically independent of the choice of origin for the coordinate system. We say that these quantities are translation and rotation invariant. One of the most successful approaches is to represent protein structures as point clouds. The most common approach is to define rotation equivariant convolution operations based on spherical harmonic as introduced by the Tensor Field Networks [22] (TFN). The Cormorant networks [23] further improved this idea with Clebsch-Gordan non-linearity enhancing the degrees of freedom of the model. TFN architectures have been successful to predict the quality of generated protein-protein complexes [24]. Instead of using spherical harmonics, a simpler approach is to define operations applied directly to vectors respecting rotation equivariance. The geometric vector perceptron [25] (GVP) uses linear operations to compose vector features with gating [26]. Graph neural networks have been extended to equivariant graph neural networks [27]. The introduction of transformers showed that recurrent neural networks are not necessary to process variable size input: attention is all you need [28]. Transformers are heavily used in natural language processing [29]. The extension of the TFN with an attention mechanism on the message passing leads to the SE(3)-Transformers [30]. It enables the modulation of angular information through spherical harmonics. Many successful methods combine transformers and geometric deep learning. The major breakthroughs come from the field of protein folding. AlphaFold [31] integrates attention in the Evoformer blocks and the structure module. The third track of the RoseTTAFold [32] model uses a SE(3)-Transformer to refine the atom coordinates during folding. The recurrent geometric network [33] (RGN2) leverages the Frenet-Serret formulas to represent the backbone of proteins. Multiple machine learning-based protein-protein interaction site prediction methods have been developed.

We introduce here PeSTo, a new parameter-free geometrical transformer that acts directly on the atoms of a protein structure. As shown in **Figure 1** and detailed in Methods, the structure is represented as a cloud of point atoms, and its geometry is described through pairwise distances and relative displacement vectors which guarantee translation invariance. The atoms are described using only their elemental names and do not require any explicit numerical parametrization such as mass, radius, charge, hydrophobicity, etc. We define a geometrical transformer operation acting on this point cloud. This operation updates the states of each atom using the states in its local neighborhood. It acts on a center atom defined with a scalar state (*q*) and a vector state (***p***) as shown in **Figure 1a**. The context of a center atom contains the neighbor’s states (*q*_*nn*_) and the neighbor’s vector states (***p***_*nn*_). The geometry of the context around a specific atom is described with the neighbors distances (*d*_*nn*_) and the neighbors normalized relative displacement (***r***_*nn*_). The interactions are encoded to create keys and values for both the scalar and vectorial states of the edges. The query is encoded from the central atom state and used with the keys and values to compute the output state with a multi-head attention operation, see **Supplementary Algorithm 1**. The geometric transformer operation can be applied to all atoms in a structure to update the state of a structure as shown in **Figure 1b**. It is translation-invariant, rotation-equivariant, and independent of the order of the atoms and order of the interactions. The attention operation allows for a dynamic number of nearest neighbors (*nn*). However, in practice, the operation is much more computationally efficient with a fixed number of nearest neighbors. In the same fashion as applying convolution operations on an image, chaining geometric transformers can propagate information at a longer range than the local context of a single operation. The main architecture is based on a bottom-up approach, starting from a small context of 8 nearest neighbors (≈ 3.4 Å radius) up to long-range interactions with 64 nearest neighbors (≈ 8.2 Å radius), see **Figure 1c**. The size of the context gradually increases allowing the model to progressively include more information while remaining cheaper in computation requirements and memory for deep models. The residual connection between geometric transformers enables us to train deeper neural network architectures. Two additional modules aggregate the atom-based geometric description at the residue level independently of the number of atoms within a residue (geometric residue pool) and predict whether each protein is at an interface or not (interface model).

**Figure 1.**
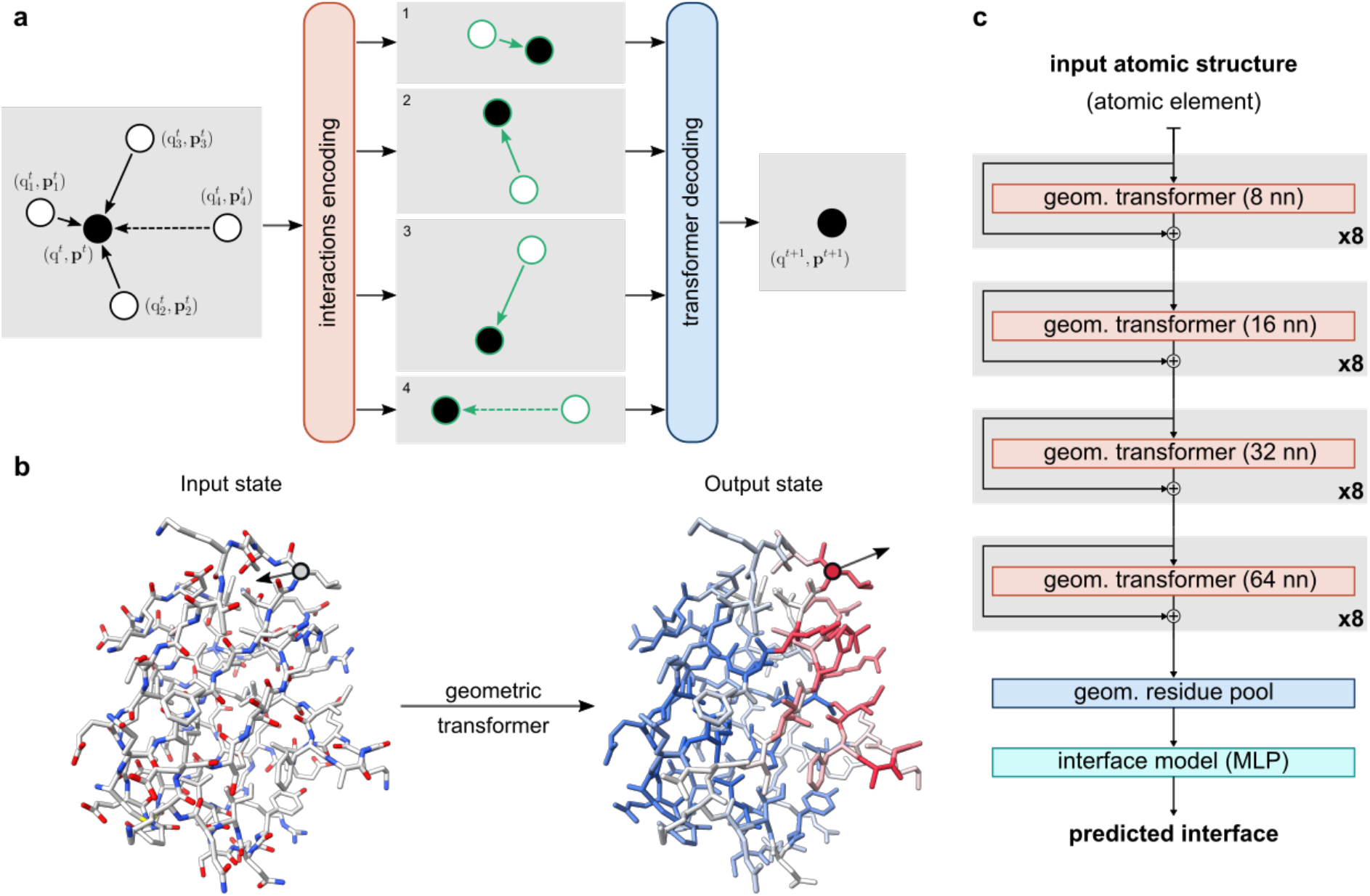
Overview of the PeSTo method. (**a**) Primary geometric transformer acting on the scalar and vectorial state of an atom at layer *t*. The interactions between the central atom and the nearest neighbors are encoded. A transformer is used to decode and filter the interactions information and to compute the new state of the central atom. (**b**) Example of application of the primary geometric transformer to all atoms in a structure. (**c**) The architecture of the PeSTo for the prediction of interactions interfaces. The model is composed of multiple layers of geometric transformers with a set number of nearest neighbors (nn) and residual connections. The structure is reduced to a residue representation through a transformer-based geometric pooling. The residue states are collapsed, and the final prediction is computed from a multi-layer perceptron (MLP).

## Results

### Protein-protein interface prediction

We trained with PDB data (see Methods) a PeSTo model to predict which residues make part of a protein interface as flagged by an output ranging from 0 to 1 (**Figure 2a**). Zero values mean that the residue is predicted to not be engaged in interactions, while with 1 values it is predicted to be at an interface. In practice, the actual value of the prediction reflects the confidence of the prediction at the residue level, with values farther from 0.5 implying higher confidence, see **Supplementary Figure 1**.

**Figure 2.**
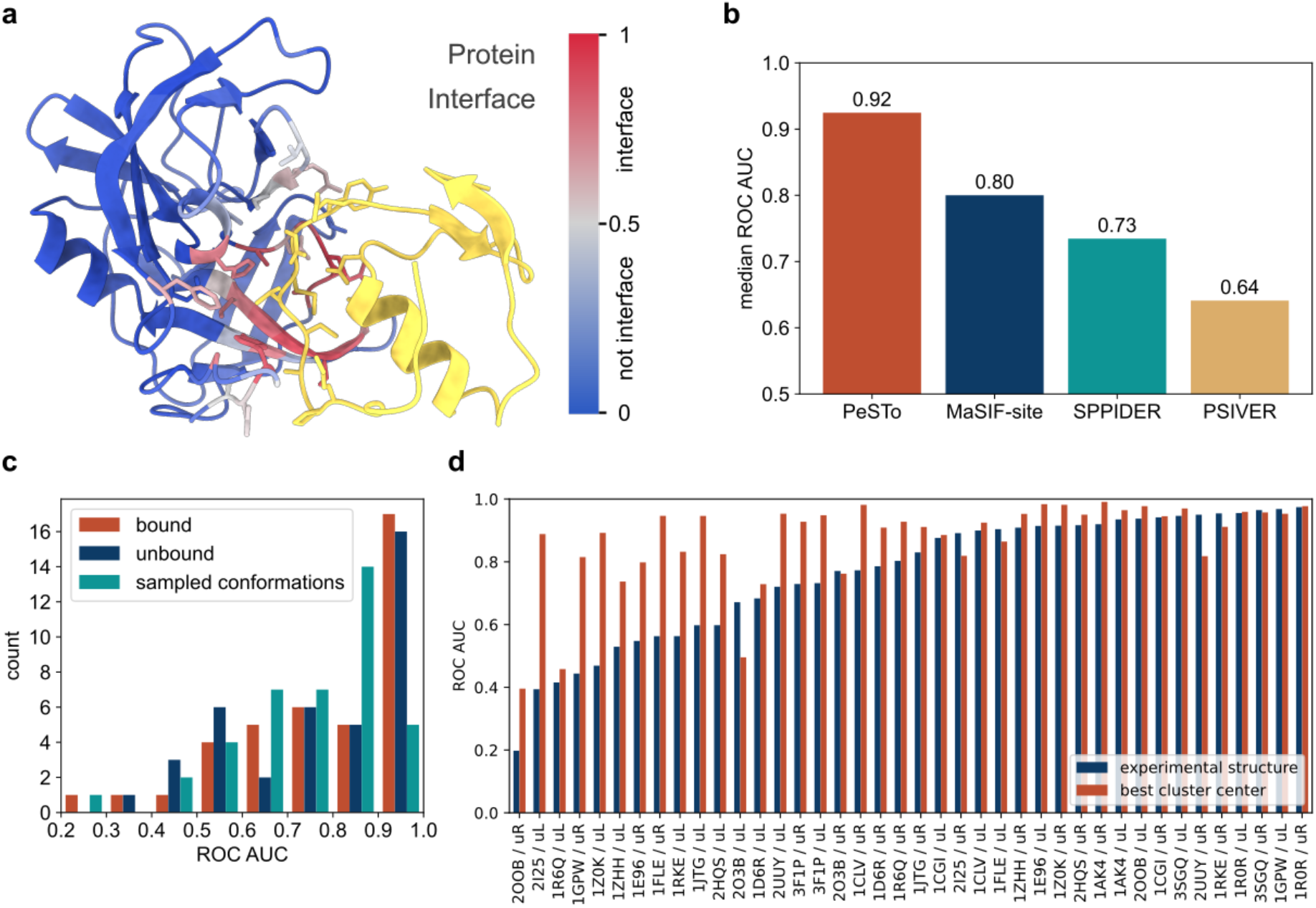
Assessment of protein-protein interface predictions with PeSTo. (**a**) Example of protein-protein interface prediction for the unbound conformation of Streptogrisin B (PDB ID: 2QA9) as can be retrieved at http://pesto.epfl.ch. The confidence of the predictions is represented with a gradient of color from blue for non-interfaces to red for interfaces. The ligand in yellow was subsequently added based on the structure of the complex (PDB ID: 3SGQ) to show the quality of the prediction. (**b**) Comparison against other methods for protein-protein interface prediction. The methods are evaluated on our groundtruth. (**c**) Benchmark of our method on bound and unbound experimental structures, as well as the average ROC AUC on 1000 sampled conformations from 1 µs MD simulations for 80 subunits taken from the PPDB5 dataset. (**d**) Clustering of predicted interfaces on 1 µs MD simulations compared against the predicted interface of the experimental structure for the unbound receptor (uR) and ligand (uL) for 20 complexes taken from the PPDB5 dataset.

On the dataset of 53 structures used to benchmark MaSIF-site (one of the best algorithms currently available), which we excluded from our training set at 30% sequence similarity, PeSTo reaches a median receiving operating characteristic (ROC) area under the curve (AUC) of 0.92 against 0.8 for MaSIF-site followed by SPPIDER [16] and PSIVER [34] (**Figure 2b**). The interfaces predicted by PeSTo have a higher ROC AUC than all other methods benchmarked here for 38 out of 53 structures.

To further showcase the quality of the predictions in real-world applications, we tested proteins from the Protein-Protein Docking Benchmark 5.0 [35] (PPDB5) dataset in their unbound conformations. The example in **Figure 2a** indeed shows PeSTo recovering the interaction interface of Streptogrisin B with ovomucoid from its unbound conformation (0.93 Å RMSD) with a ROC AUC of 96%. The short time needed to run the model (300 ms for a 100 kDa protein from PDB load to prediction on a single GPU, details in **Supplementary Figure 3**) allows us to evaluate snapshots from large structural ensembles efficiently, even datasets extracted from molecular dynamics (MD) simulations. We applied PeSTo for protein-protein interface prediction on 1000 frames from eighty 1 µs-long atomistic MD simulations of the unbound and bound experimentally derived conformations of subunits taken from the Protein Docking Benchmark version 5 (**Figure 2c**). The experimentally determined bound and unbound along with the computationally sampled conformations have a median ROC AUC of 85%, 81%, and 79%, respectively. Therefore, the model performs almost as well on experimentally solved bound and unbound conformations. Although overall the ROC AUC decreases with a higher RMSD from the bound structure (**Supplementary Figure 3**) our method is still able to recover the interface with a ROC AUC higher than 80% for most structures and MD-sampled conformations.

In some cases, processing full MD trajectories of unbound proteins with PeSTo identifies certain interfaces better than when PeSTo is run on the starting structures, which suggests a major practical application of our method to real-life situations (**Figure 2d**). Out of 40 cases, the model correctly recovers 21 interfaces (ROC AUC above 0.8) from the unbound conformation, independently of the MD sampling and clustering. Out of the remaining 19 cases, we observe a recovery of the interacting interface from an unbound conformation for 12 subunits using MD to sample conformations and additional clustering of the interfaces.

For instance, PeSTo predicts no interface for the experimentally solved structure of the unbound porcine pancreatic elastase (PDB ID 9EST), with a ROC AUC score of only 56%. The unbound experimental conformation has an RMSD of 1.2 Å from the bound complex with elafin (PDB ID 1FLE). However, MD simulation starting from the unbound porcine pancreatic elastase alone shows a conformation switch leading to the recovery of the interaction interface with elafin with a cluster center ROC AUC of 92%. Inspecting the simulation unveils that the motion of a loop in elastase is required to allow elafin to enter the pocket and accommodate an inter-molecular β- sheet that stabilizes the complex as solved experimentally (**Supplementary Figure 4**).

### General protein interface prediction

In light of the results for protein-protein interface predictions, we generalized the model to find and identify more types of interfaces, resulting in a generalized PeSTo model that predicts protein interaction interfaces with other proteins, DNA/RNA, ions, ligands, and lipids. We trained a generalized PeSTo model with PDB structures featuring all the kinds of expected interactions, as described in Methods. The interface predictions for protein-DNA/RNA interfaces are almost as good as for protein-protein interfaces, reaching ROC AUC of 0.89 for a validation set (**Figure 3a**). The generalized model can also detect ion, ligand, and lipid interfaces with ROC AUCs of 0.88, 0.84, and 0.77, respectively on each validation set. The model does experience some confusion between ions and ligands as revealed by the confusion matrix (**Supplementary Figure 5**).

**Figure 3.**
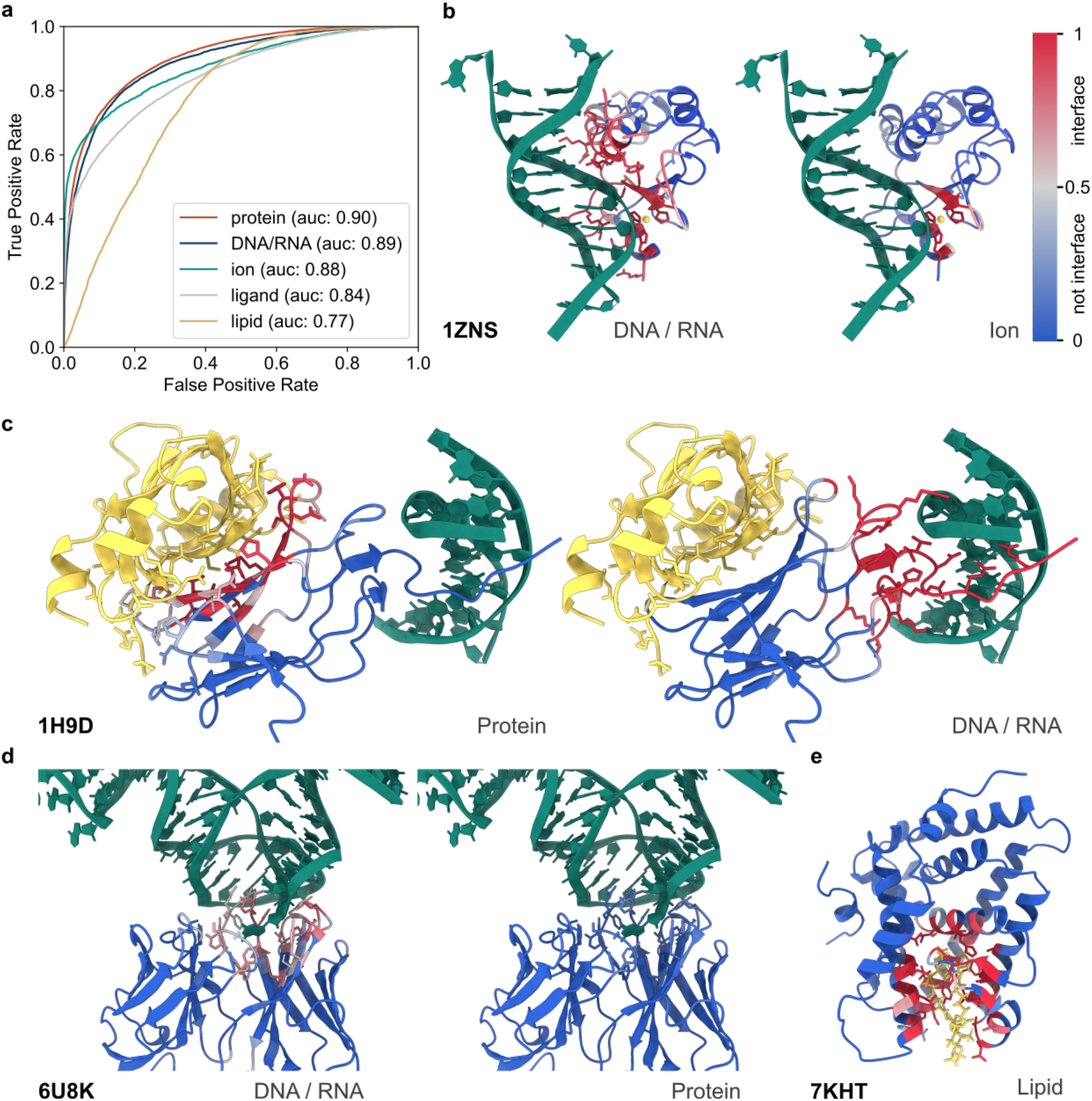
General protein interaction interface prediction with PeSTo. (**a**) ROC curve for the predictions of different types of interfaces with PeSTo. (**b**,**c**,**d**,**e**) Example of predicted interfaces. The confidence of the predictions is represented with a gradient of color from blue for non-interfaces to red for interfaces. The structures in yellow and green were added subsequently from the reference complexes. (**b**) Colicin E7 endonuclease domain in complex with DNA and a zinc ion (PDB ID: 1ZNS). (**c**) core-biding factor subunit alpha-2 in complex with core-binding factor subunit beta and DNA (PDB ID: 1H9D). (**d**) Antigen-binding fragment in complex with RNA (PDB ID: 6U8K). (**e**) Steroidogenic factor 1 bound to a phosphoinositide (PDB ID: 7KHT).

We next illustrate the generalized PeSTo with five examples that attest to its capacity to discern among various interfaces, even when they are overlapping or under-represented in the PDB. The first example (**Figure 3b**) corresponds to the colicin E7 endonuclease domain, which binds DNA through an interface that includes a Zn(II) ion (PDB ID 1ZNS). Running the apo-protein through the generalized PeSTo returns correct predictions for both interfaces, even in the overlapping part. The second case (**Figure 3c**) corresponds to the complex formed by RUNX1 with a dsDNA bound to one end and the protein CBFβ bound to the other (PDB ID 1H9D). Running the isolated RUNX1 through the generalized model returns clear, accurate interfaces through the DNA and protein channels. In the third example (**Figure 3d**) we challenge the generalized model with the structure of an antibody that binds RNA (PDB ID 6U8K) as opposed to most crystallized antibodies which are bound to other proteins. The generalized model correctly predicts no interface for proteins and the correct interface for RNA.

On interfaces with lipids, the generalized PeSTo performs less well, probably because of limited training data (only 0.7% of the utilizable data we compiled). However, in practice, we observe that the model is able to accurately detect lipid-binding pockets for soluble proteins (exemplified by the steroidogenic factor in **Figure 3e**) and even the membrane-spanning region of transmembrane proteins (**Supplementary Figure 6**), despite not specifically trained for any of these, in both cases being able to detect specific pockets for lipids as stronger scores. Many interfaces with lipids are only partially resolved experimentally resulting in low data quality leading to an artificial drop of the ROC AUC.

## Discussion

We showed here that a geometrical transformation of protein atomic coordinates suffices to detect and classify protein interfaces at high resolution, surpassing the prediction capabilities of other methods without even the need of explicitly describing the physics and chemistry of the system, hence without the overhead of pre-computing molecular surfaces and their properties. All this with modest computational resources and at a very high speed enables the analysis of large ensembles, for example from MD simulations as presented, or large datasets like those being created by the latest generations of tertiary structure prediction tools.

To make PeSTo-based predictions for proteins available to the community, we set up a web server at https://pesto.epfl.ch/, accessible free of charge without registration. The server takes protein structures/models in PDB format and returns them right away with the B-factor column filled with the per-residue predictions. This output file can be downloaded or visualized right within the website.

Provided sufficient training data are available, the method is reusable for specific applications. For example, one could train a model to specifically identify certain kinds of interfaces or pockets. More broadly, the parameter-free PeSTo architecture is general enough that could be easily accommodated to pursue other structure-based problems such as docking or modeling interactions with materials. The description is totally agnostic to the exact physicochemical properties of the atoms in the structure, thus easily extendable to other materials and fields, and is probably also less sensitive to problems related to the starting structures such as missing atoms as compared to methods that require intermediate calculations of surface and volume. Given the ever-growing accumulation of structural information and rapid expansion of predicted foldome data, PeSTo stands as an accurate, flexible, fast, and user-friendly solution to dissect the vast and dynamic protein interaction landscape.

## Methods

### Datasets

The dataset is composed of all the biological assemblies from the Protein Data Bank [36]. The subunits are clustered using a maximum of 30% sequence identity between clusters. The clusters of subunits are grouped into approximately 70% training set (381068 chains), 15% testing set (98520 chains), and 15% validation set (87685 chains). The validation set is composed of the clusters containing any of the 53 subunits from the MaSIF-site benchmark dataset or 230 structures from the Protein-Protein Docking Benchmark 5.0 [35] (PPDB5) dataset. All the examples selected to assess the quality of the predictions from the model belong to the validation set.

### Structures processing

All models of the structure are loaded as a single structure. The chain name is tagged with the model identifier to distinguish subunits from different models. Moreover, the chain name of all non-polymer chemical molecules is tagged to have them in separate subunits. Duplicated subunits, molecules, and ions generated when concatenating multiple models are removed. The first alternate location of the atoms is kept. Water, heavy water, hydrogen, and deuterium atoms are removed from the structures.

### Features and labels

We identified the 30 most common atomic elements on PDB. The element is used as the only feature as a one-hot encoding. The input vectorial features are set to zero. The distances matrices and normalized displacement vector matrices are used as geometrical features. Amino acids, nucleic acid, ions, ligands, and lipids are selected from a list of 20, 8, 16, 31, and 4 most common molecules, respectively. Non-native molecules used to help to resolve the structure are ignored. An interface is defined as a residue-residue contact within 5 Å. All protein-protein interfaces as well as protein-DNA/RNA, protein-ion, protein-ligand, and protein-lipids interfaces are identified. The details of the interface for each subunit are stored in the dataset as an interactions types matrix (79 × 79). It enables the selection of specific interfaces as labels at the start of the training session without having to rebuild the whole dataset. The interfaces targets can be selected from any combinations of subsets from the 79 molecules available.

### Deep learning architecture

For structures with a number of atoms smaller than the set number of nearest neighbors (nn), the additional non-existent interactions are sent to a sink node with a scalar and vector state set to zero. The input features are embedded to an input state size of S = 32 with a 3 layers neural network. Multiple geometric transformers (N_head_ = 2, D_key_ = 3) can be put together to create a deep model. The geometric residue pool module aggregates the encoding at the atomic level of the structure to a residue description by using a self-attention operation with 4 heads. The second module is a multi-layer perceptron decoding the state of all residues and computing the prediction.

### Training

Two models based on the same architecture were trained. The first one predicts protein-protein interface and the second one is more general and predicts protein interfaces with protein, DNA/RNA, ligand, ion, or lipid. The best neural networks architecture was trained for 4 days on a single NVIDIA V100 (32 GB) GPU. Subunits with a maximum of 8192 atoms (≈ 100 kDa) without hydrogens are used to limit the memory requirement during training. Subunits with less than 48 amino acids are ignored during training. The effective protein-protein interfaces dataset is composed of 86374 subunits for training and 21587 subunits for testing. The effective generalized protein interfaces dataset is composed of 113805 subunits for training and 29786 subunits for testing.

### Evaluation

Our method was compared with MaSIF-site [7,17], SPPIDER [16], and PSIVER [34]. MaSIF-site is the best available method for protein-protein interface prediction. SPPIDER is a long-standing and well-tested method used as a reference for protein-protein interface prediction. PSIVER only uses sequence information and is benchmarked to show the difference in performance between structure-based and sequence-based methods.

### Molecular dynamics

20 complexes from the PPDB5 dataset were selected based on the resolution of the structure and the difficulty to parametrize. For each, we performed a 1 µs simulation of the subunits alone for the bound receptor (bR), unbound receptor (uR), bound ligand (bL), and unbound ligand (uL). 500 frames per simulation are used to evaluate the model for a total of 400 000 frames.

## Acknowledgments

We thank the Swiss National Science Foundation (SNSF) for support (grant number 205321_192371 to M.D.P) and the Swiss National Supercomputing Center (CSCS) for the generous time allocations used to run molecular dynamics simulations.

## Supplementary information

### Algorithm 1

Geometric transformer

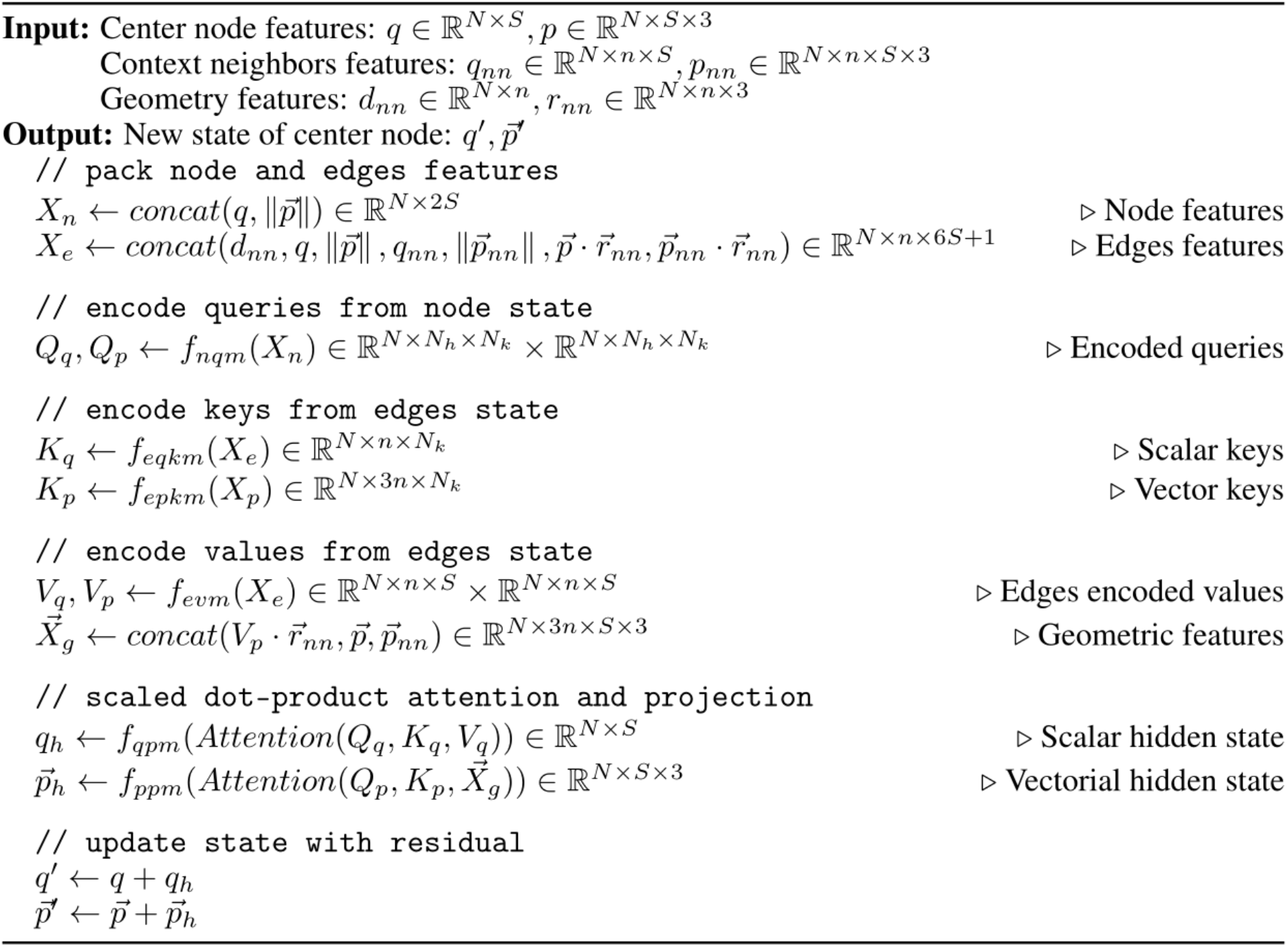

**Supplementary Algorithm 1** | **Geometric transformer**. Each geometric transformer is composed of 5 neural networks of 3 layers with an exponential linear unit (ELU) activation function. The hidden layers width corresponds to the state size (S). The multi-layers perceptrons (MLP) are the node query model (f_nqm_), encoding scalar key model (f_eqkm_), encoding vector key model (f_epkm_), encoding value model (f_evm_), and scalar state projection model (f_qpm_). The vectorial hidden state is projected with a weighted sum (f_ppm_) in order to preserve the rotation equivariance of the operation. The output vector state belongs to the span of the geometry and vector states.

**Supplementary Figure 1.**
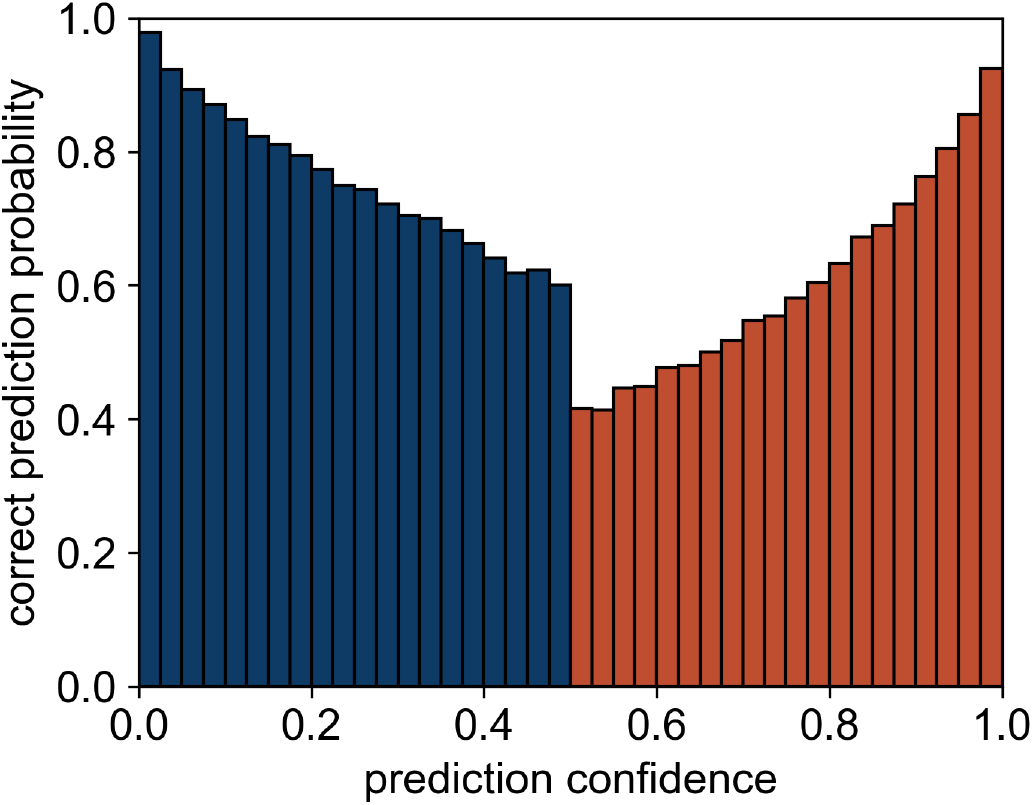
Prediction quality estimation. Estimated correlation between protein-protein interface prediction confidence and prediction quality. Evaluated on 8192 structures randomly sampled from the validation dataset.

**Supplementary Figure 2.**
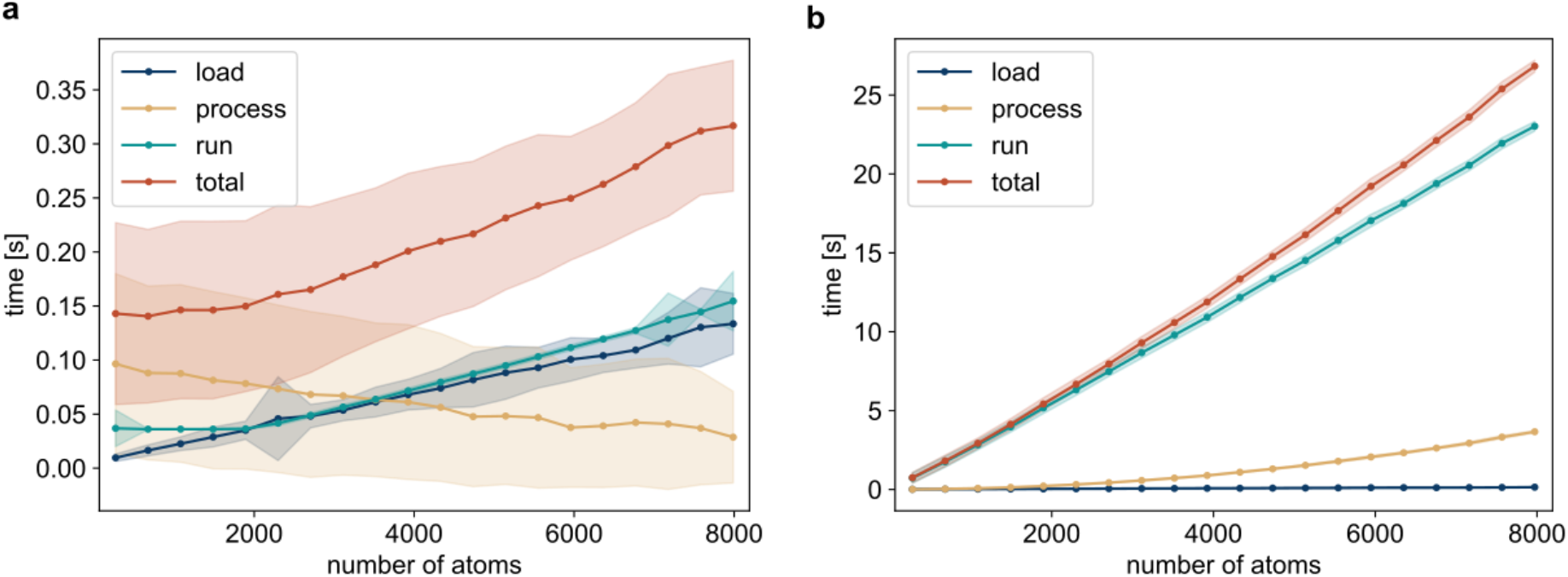
Profiling of the run time of PeSTo as a function of the size of the structure. Run time evaluated (**a**) on GPU (NVIDIA RTX 2080 Ti) and (**b**) on CPU only (Intel i9-9900K). For structures of around 100 kDa (8000 atoms), the average total runtime is 300 ms with 130 ms to parse the file, 30ms to process the structure and 140 ms to run the inference on a high-end GPU.

**Supplementary Figure 3.**
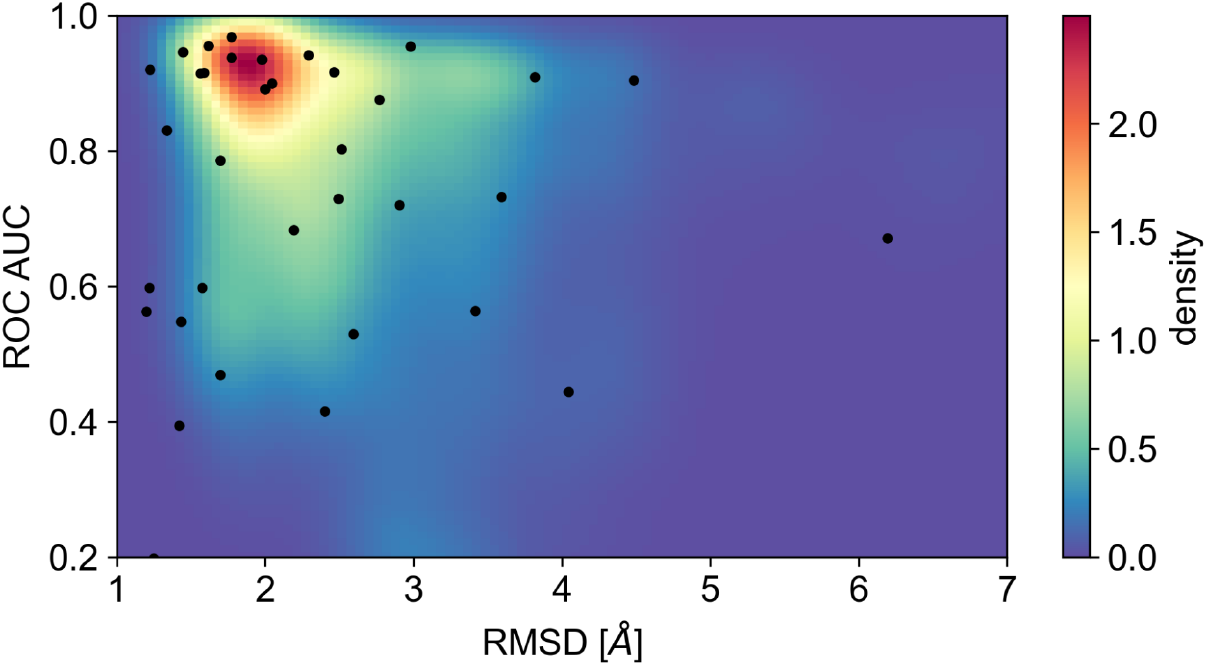
ROC AUC as a function of RMSD for different conformations for the 80 simulated subunits from the PPDB5 dataset. The RMSD is computed from the bound conformation of the subunits in the reference complex. Starting conformations are indicated with a black dot.

**Supplementary Figure 4.**
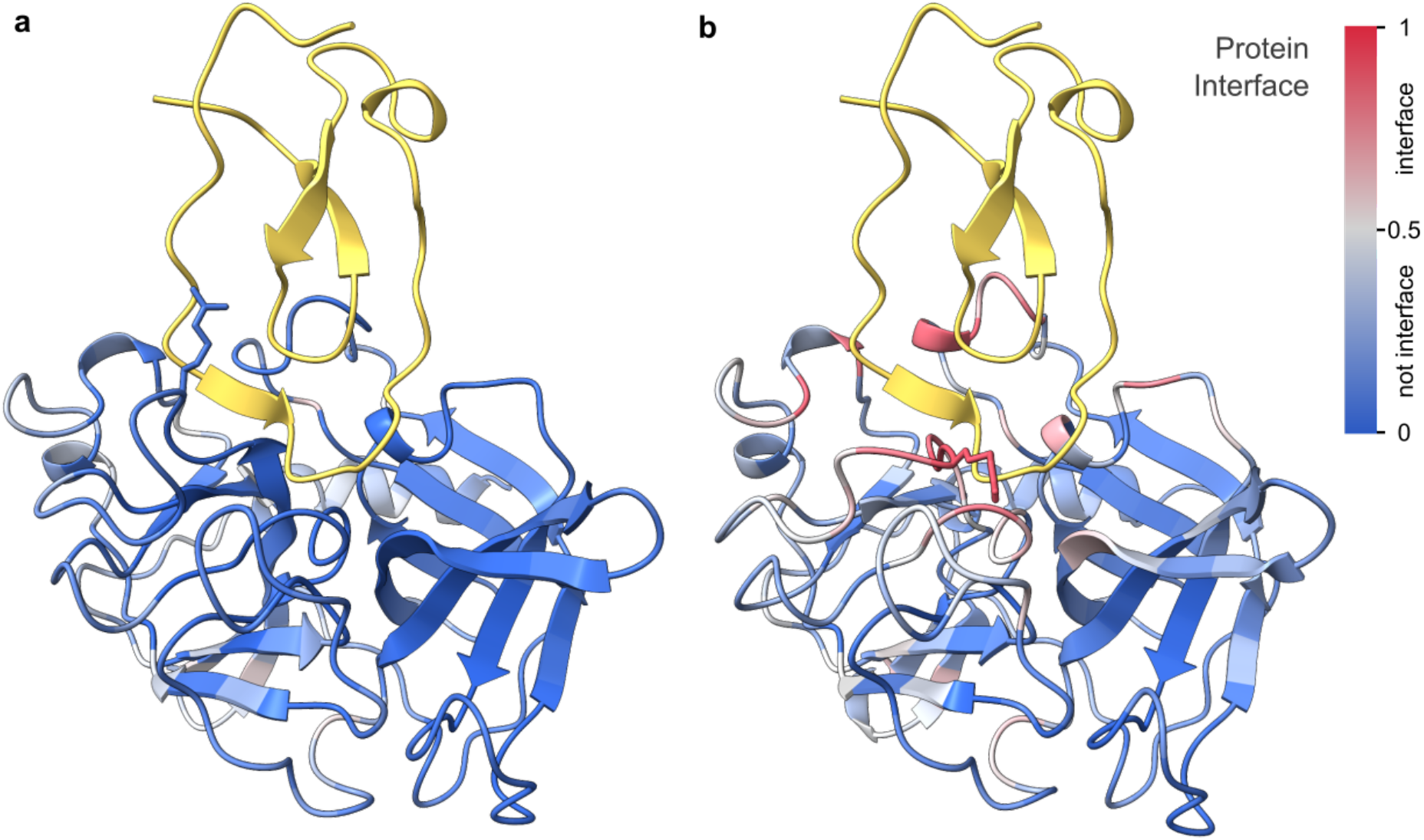
Example of recovered interaction interface from conformation change. The confidence of the predictions is represented with a gradient of color from blue for non-interfaces to red for interfaces. The structure in yellow was added subsequently from the reference complexes (PDB ID 1FLE). (**a**) Protein-protein interface prediction on the experimentally resolved structure of unbound porcine pancreatic elastase (PDB ID 9EST). (**b**) Protein-protein interface prediction on an open conformation sampled using MD and selected using clustering on the conformations. Arg217A is shown in licorice to illustrate the rearrangement of the loop region containing Arg217A opening up the interface.

**Supplementary Figure 5.**
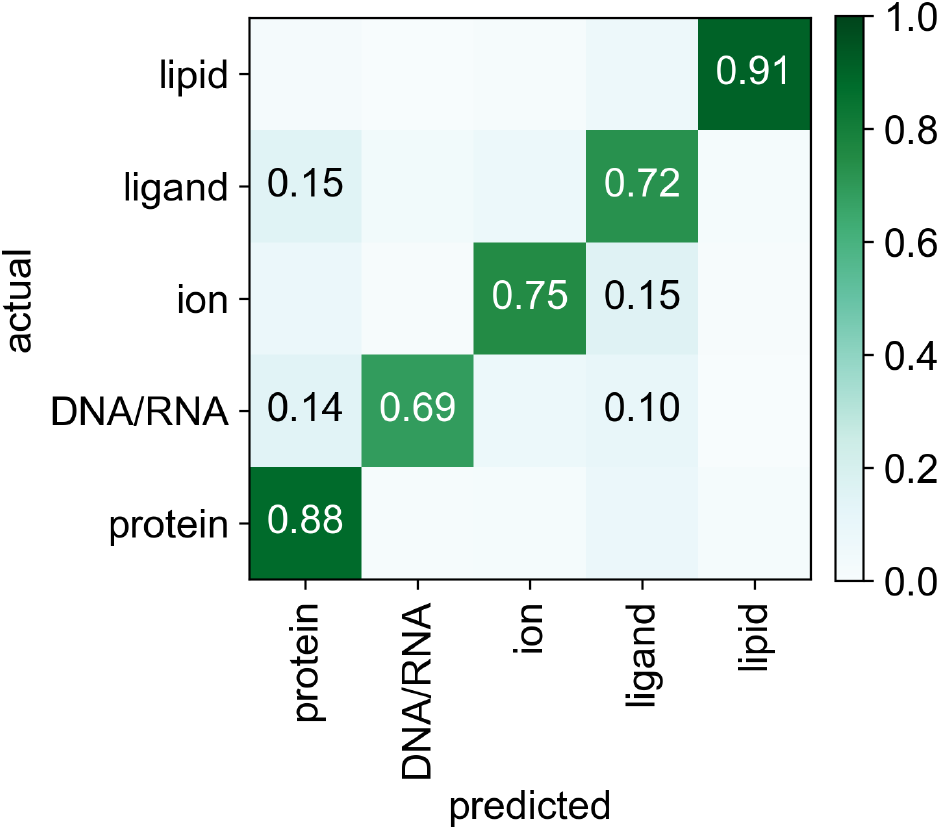
Confusion matrix between actual and predicted interfaces at the residue level. Each interface type is randomly sampled equally at ∼3600 residues per class. The confusion matrix is normalized per actual interface. We observed nucleoside triphosphates (ATP, GTP) and diphosphates (ADP, GDP) pockets misclassified as DNA/RNA binding regions, a reasonable confusion given the chemical similarity of all these molecules.

**Supplementary Figure 6.**
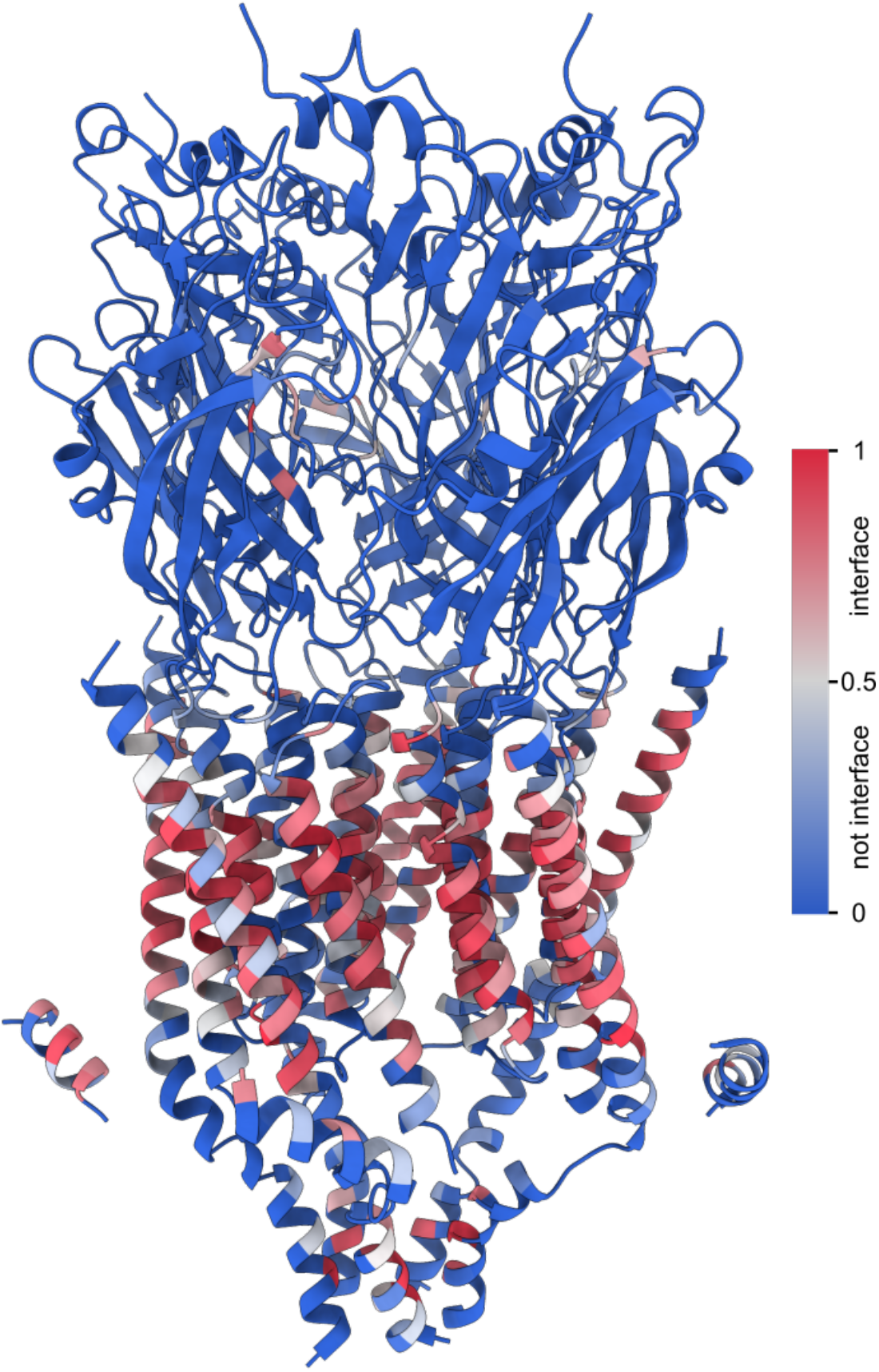
Example of lipid interface prediction for transmembrane protein. Homopentameric 5-HT 3A serotonin receptor (PDB ID: 6Y5B).

